# Objectively measured physical activity levels and sedentary time in children and adolescents with sickle cell anemia

**DOI:** 10.1101/325969

**Authors:** Hugo Nivaldo Melo, Simone Joanna-Maria Stoots, Marijn Aimee Pool, Vitor Oliveira Carvalho, Max Luan De Carvalho Aragão, Ricardo Queiroz Gurgel, Charles Agyemang, Rosana Cipolotti

## Abstract

The aim of this study was to identify the levels of physical activity and sedentary behaviour of children and adolescents with sickle cell disease (SCA) compared to healthy individuals. A cross-sectional study with a quantitative approach was performed at a reference center for the treatment of patients with hemoglobinopathies in northeastern Brazil. Patients were recruited between October 2015 and January 2017. Eligible participants answered a Physical Activity Questionnaire for Older Children and Adolescents (PAQ-C) and were instructed to use an ActiGraph wGT3X-BT triaxial accelerometer for seven consecutive days. The analysis of the results was performed using the SPSS software (version 13.0). Differences between means were analysed using the Mann-Whitney U test and the chi-square test was used to evaluate the proportions of occurrence of categorical variables, comparing patient and controls groups. Among the 352 patients in the follow-up, 64 met the inclusion criteria and agreed to participate. Of those, 14 did not use the accelerometer during the 7 consecutive days and were excluded. 50 patients (and their 50 controls) were then evaluated. We observed a statistically significant difference between cases and controls in the variables “total time of moderate and vigorous physical activity” (p=0.009 and p=0.0001, respectively) and “daily mean of moderate and vigorous physical activity” (p=0.005 and p=0.003). There was also a significant difference among cases and controls in the following variables: “metabolic equivalent” (MET), with p=0.04, total of steps (p=0.04) and “total caloric expenditure” (p=0.0001), with the worst performances for the group of patients with SCA. Children and adolescents with SCA presented lower levels of physical activity than healthy children and adolescents, both when evaluated by the PAQs or by the accelerometer. The results suggest the need to develop specific programs aimed at promoting physical activity levels and reducing sedentary behaviour among young individuals with SCA.

## Author Summary

Sickle cell anemia is a hereditary and hematological disease that occurs due to the abnormal production of red blood cells. On the other hand, childhood physical activity has beneficial effects on health, both in the short and long terms, and may reduce risk factors for chronic diseases. The aim of this study was to identify the levels of physical activity and sedentary behaviour of children and adolescents with sickle cell disease compared to healthy individuals. Participants answered a Physical Activity Questionnaire for Older Children and Adolescents (PAQ-C) and were instructed to use an ActiGraph wGT3X-BT triaxial accelerometer for seven consecutive days. We observed children and adolescents with sickle cell anemia presented lower levels of physical activity than healthy children and adolescents, both when evaluated by the PAQs or by the accelerometer. The results suggest the need to develop specific programs aimed at promoting physical activity levels and reducing sedentary behaviour among young individuals with sickle cell anemia.

## Introduction

Sickle cell anemia (SCA) is a neglected tropical disease [1] characterized by a point mutation in the β-chain hemoglobin (Hb) gene. When deoxygenated, HbS, the Hb resulting from the mutation, polymerises, resulting in a change in the red blood cells and assuming a sickle shape. The falcized red blood cells can obstruct the microcirculation, resulting in ischemia-reperfusion injury, with inflammatory cytokines, pain and functional impairment, particularly in the musculoskeletal system [2], which can lead to defensive sedentary behaviour even among young patients.

Previously, it has been demonstrated that childhood physical activity (PA) has beneficial effects on health, both in the short and long terms, and may reduce risk factors for chronic diseases [3]. More recently, one study concluded that moderate physical exercise is not harmful for patients with SCA [4], and, in an animal model (rats with SCA), it is suggested that PA could be beneficial in the clinical course of the disease [5].

The evaluation of PA is complex due to its multi-dimensional characteristics. Questionnaires that use semi-quantitative scales, such as the Physical Activity Questionnaire for Older Children and Adolescents (PAQ-C), have the advantage of easier applicability, but they are influenced by the interviewee’s perception of their level of physical activity. Thus, objective methods are preferable and the accelerometer is an instrument that generates quantitative results and thus confers objectivity to the assessment of BP in children and adolescents [6].

Thus, this study aimed to evaluate the level of PA and sedentary behaviour of children and adolescents with SCA compared to healthy individuals.

## Methods

### Design

This is a cross-sectional study carried out at the outpatient clinic of a university center in the northeast of Brazil, which is a regional reference for the treatment of patients with SCD. After initial screening, eligible patients and healthy controls completed the Physical Activity Questionnaire for Older Children and Adolescents (PAQ-C) [7,8]. Subsequently, participants and their caregivers were instructed on how to properly use the ActiGraph wGT3X-BT triaxial accelerometer and to use it for seven consecutive days.

### Population

Patients were recruited between October 2015 and January 2017. Among the patients with SCA (homozygous for HbS) confirmed by hemoglobin electrophoresis, individuals who were between 6 and 18 years of age and in a stable clinical condition were considered eligible, if they had not received blood transfusions in the last three months and if they were without acute complications for at least one month, before being included in this study. Patients with neurological or orthopedic impairment were excluded.

The control group consisted of healthy children and adolescents recruited at a local public school and matched for age and sex with the patients.

### Ethics Statement

This study was approved by the Research Ethics Committee involving Human Beings of the Federal University of Sergipe (protocol: 30661314.0.0000.5546). All guardians responsible for the patients and controls signed a free and informed consent form.

### Laboratory tests

The results of hematological examinations (hemoglobin, hematocrit, erythrocyte counts, platelets, leukocytes, neutrophils, reticulocytes, indirect bilirubin, mean corpuscular volume, lactate dehydrogenase) and hemoglobin electrophoresis (fetal hemoglobin and hemoglobin S) were obtained retrospectively from a database especially created for this research. Laboratory tests were performed under stable clinical conditions within four weeks prior to the application of the accelerometer. All the exams were performed at the Central Laboratory located at the service itself, using the same standardised techniques and equipment.

### Medication use

All patients were using folic acid supplements (2 mg/day). Patients on hydroxyurea received an initial dose of 15 mg/kg/day and were currently using the standard dose (20 to 35 mg/kg/day) for at least 12 months [9].

### Physical activity questionnaire for older children and adolescents (PAQ-C)

All patients included in this study completed the Brazilian version of PAQ-C [8], composed of nine questions about sports, games, and other physical activities at school and at leisure activities. This questionnaire aims to provide a complete picture of the type and amount of PA that the participant had been performing in the last seven days prior to the interview. Each question was scored on a scale of 1 to 5, being: very sedentary (1), sedentary (2), moderately active (3), active (4) or very active (5). In order to determine the final score, the mean of all responses was calculated.

### PA measurements

The ActiGraph GT3X Accelerometer (ActiGraph LLC, Pensacola, FL, USA) was used to objectively monitor the time spent in PA and sedentary behaviour. The accelerometer was worn on an elastic belt and participants were instructed to position it on the hip line of their dominant side. Participants used the device for 7 consecutive days, including two weekend days for at least 10 hours a day [10,11]. The study team instructed and monitored the children and their caregivers to remove the monitor during aquatic activities and during sleep. The accelerometer was initialised by the researcher responsible for the study through the manufacturer’s software (ActiLife version 6). In order to record the movement in counts per minute, the count was set to 60 second epochs. The time of sedentary behaviour was defined by <100 count per minute [12,13].

Values between 100 and 1999 counts per minute were recorded as light PA (LPA) [13]. The time spent on moderate PA (MPA) and vigorous PA (VPA) was calculated based on cutoffs of 2000 and 4000 counts per minute, respectively [14]. The PA of each individual was categorised in the three intensity levels (LPA, MPA and VPA) and the average daily sedentary time had been recorded. The time spent in moderate/vigorous PA (MVPA) was calculated as the sum of MPA and VPA.

The daily percentage of all PA intensity levels was calculated based on the time spent at each intensity level, including sedentary time [15]. For comparison, the children were considered to be in accordance with the recommendations of the AP when the mean MPVA over all measured days was 60 min or more [16]. The mean time measured for both weekdays and weekends was calculated by summarising the sedentary time and the time spent at different BP intensities.

### Statistical analysis

The data analysis was performed using SPSS version 13.0 for Windows
(SPSS, Inc., Chicago, IL, USA). Quantitative variables were described as means and standard deviations. All variables were checked for normality prior to analysis using the Kolmogorov-Smirnov test. Differences between means were analysed using the Mann-Whitney U test and the chi-square test was used to evaluate the proportions of occurrence of categorical variables, comparing patient and control groups. Differences in time spent at different PA intensities and the mean times measured on weekdays and weekends were analysed by using the t-test for paired samples. The chi-square test was used to determine differences in the percentage of time spent at different PA intensities. The significance level used was 5% (p<0.05).

## Results

We accessed a registry of patients with SCA who regularly attended the outpatients’ clinic in the study institution (352 children and adolescents). 288 patients did not meet the inclusion criteria. 64 patients were considered eligible, but 14 were later excluded because they did not use the accelerometer for seven consecutive days. Thus, 50 patients were included in this study, of which 60% were male with a mean age of 12.02 ± 3.6 years.

The control group was selected from a public school located in the same city. The students were asked to participate with consent of their guardians and, if they did not have any relatives diagnosed with SCA and if they had no chronic disease nor acute disease during the days of using the accelerometer, were paired to patients with SCA according to sex and age.

**Table 1:**
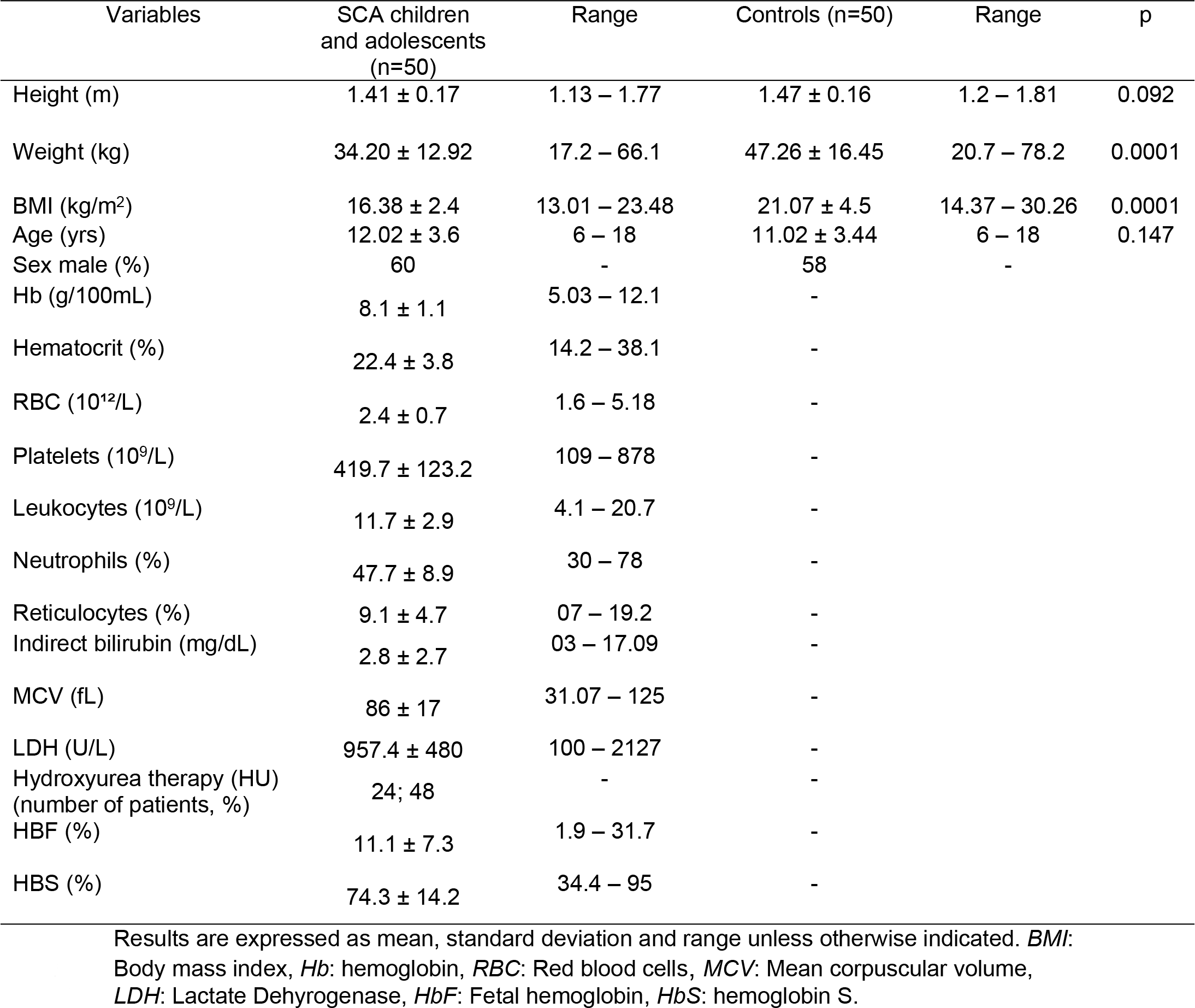
Characterization of study participants

| Variables | SCA children and adolescents (n=50) | Range | Controls (n=50) | Range | p |
| --- | --- | --- | --- | --- | --- |
| Height (m) | 1.41 ± 0.17 | 1.13 – 1.77 | 1.47 ± 0.16 | 1.2 – 1.81 | 0.092 |
| Weight (kg) | 34.20 ± 12.92 | 17.2 – 66.1 | 47.26 ± 16.45 | 20.7 – 78.2 | 0.0001 |
| BMI (kg/m<sup>2</sup>) | 16.38 ± 2.4 | 13.01 – 23.48 | 21.07 ± 4.5 | 14.37 – 30.26 | 0.0001 |
| Age (yrs) | 12.02 ± 3.6 | 6 – 18 | 11.02 ± 3.44 | 6 – 18 | 0.147 |
| Sex male (%) | 60 | - | 58 | - |  |
| Hb (g/100mL) | 8.1 ± 1.1 | 5.03 – 12.1 | - |  |  |
| Hematocrit (%) | 22.4 ± 3.8 | 14.2 – 38.1 | - |  |  |
| RBC (10<sup>12</sup>/L) | 2.4 ± 0.7 | 1.6 – 5.18 | - |  |  |
| Platelets (10<sup>9</sup>/L) | 419.7 ± 123.2 | 109 – 878 | - |  |  |
| Leukocytes (10<sup>9</sup>/L) | 11.7 ± 2.9 | 4.1 – 20.7 | - |  |  |
| Neutrophils (%) | 47.7 ± 8.9 | 30 – 78 | - |  |  |
| Reticulocytes (%) | 9.1 ± 4.7 | 07 – 19.2 | - |  |  |
| Indirect bilirubin (mg/dL) | 2.8 ± 2.7 | 03 – 17.09 | - |  |  |
| MCV (fL) | 86 ± 17 | 31.07 – 125 | - |  |  |
| LDH (U/L) | 957.4 ± 480 | 100 – 2127 | - |  |  |
| Hydroxyurea therapy (HU) (number of patients, %) | 24; 48 | - | - |  |  |
| HbF (%) | 11.1 ± 7.3 | 1.9 – 31.7 | - |  |  |
| HbS (%) | 74.3 ± 14.2 | 34.4 – 95 | - |  |  |

Results are expressed as mean, standard deviation and range unless otherwise indicated. *BMI*: Body mass index, *Hb*: hemoglobin, *RBC*: Red blood cells, *MCV*: Mean corpuscular volume, *LDH*: Lactate Dehydrogenase, *HbF*: Fetal hemoglobin, *HbS*: hemoglobin S.

All 50 patients and 50 controls used the accelerometer for seven consecutive days without any complications. The clinical characteristics of both groups (patients with SCA and healthy controls) are described in Table 1. The groups were similar in terms of age and distribution by sex, meeting the pairing criteria, but presented a statistically significant difference in the means of body mass and body mass index (BMI), with p=0.0001.

**Table 2:** Physical Activity and Sedentary Behavior Variables

| Variables | SCA children and adolescents (n=50) | Range | Controls (n=50) | Range | p |
| --- | --- | --- | --- | --- | --- |
| Sedentary Time (min) | 2918.12 ± 848.3 | 1271 – 5208 | 2865.5 ± 914.4 | 1363 – 6682 | 0.775 |
| Sedentary Time (min/day) | 416.87 ± 106.03 | 158.87 – 651 | 409.35 ± 114.3 | 170.37 – 835.25 | 0.639 |
| Sedentary Time (%) | 56.84 ± 13.92 | 30.66 – 95.54 | 52.54 ± 11.76 | 32.71 – 77.46 | 0.453 |
| Light PA (min) | 2279.04 ± 913.51 | 740 – 5208 | 2359.84 ± 751.95 | 880 – 3517 | 0.630 |
| Light PA (min/day) | 325.57 ± 114.18 | 92.5 – 651 | 337.12 ± 93.99 | 110 – 439.62 | 0.237 |
| Light PA (%) | 42.96 ± 10.77 | 22.98 – 95.54 | 47.73 ± 14.84 | 20.14 – 59.79 | 0.424 |
| Moderate PA (min) | 134.9 ± 95.5 | 15 – 447 | 190.18 ± 110.9 | 6 – 498 | 0.009 |
| Moderate PA (min/day) | 19.27 ± 11.94 | 1.87 – 59.62 | 27.16 ± 13.87 | 0.75 – 62.25 | 0.005 |
| Moderate PA (%) | 2.66 ± 1.8 | 0.35 – 8.31 | 3.51 ± 1.93 | 0.15 – 8.3 | 0.368 |
| Vigorous PA (min) | 25.76 ± 33.05 | 0 – 202 | 54.94 ± 59.85 | 0 – 370 | 0.0001 |
| Vigorous PA (min/day) | 3.68 ± 4.13 | 0 – 25.25 | 7.84 ± 7.48 | 0 – 46.25 | 0.003 |
| Vigorous PA (%) | 0.51 ± 0.61 | 0 – 3.52 | 1.11 ± 0.93 | 0 – 5.03 | 0.351 |
| MPVA (min) | 160.66 ± 148.88 | 30 – 752 | 245.12 ± 538.05 | 20 – 1014.83 | 0.712 |
| MPVA (min/day) | 22.95 ± 18.6 | 3.7 – 94 | 35.01 ± 66.6 | 2.5 – 306.2 | 0.705 |
| MPVA (%) | 3.17 ± 4.35 | 0.47 – 9.12 | 4.62 ± 2.49 | 0.53 – 11.25 | 0.423 |
| MET | 1.71 ± 0.4 | 1.17 – 2.45 | 1.87 ± 0.39 | 1.12 – 2.57 | 0.04 |
| Current PA recommendations | 8 | - | 30 | - |  |
| Kcal total | 1015.73 ± 516.83 | 701.56 – 2659.73 | 2404.31 ± 1308.22 | 806.23 – 5178.81 | 0.0001 |
| Sitting (%) | 18.3 ± 5.17 | 7 – 32 | 19.16 ± 5.83 | 11 – 36 | 0.153 |
| Standing (%) | 24.94 ± 7.34 | 6 – 40 | 25.64 ± 7.43 | 10 – 39 | 0.682 |
| Lying (%) | 5.12 ± 3.8 | 0 – 19 | 6.24 ± 5.75 | 0 – 28 | 0.254 |
| Total steps | 51010.52 ± 19600.13 | 16057 – 120567 | 59105.40 ± 22650.89 | 20583 – 137817 | 0.04 |

Results are expressed as mean, standard deviation and range unless otherwise indicated. PA: physical activity, MPVA: moderate-to-vigorous physical activity.

There was a statistically significant difference between the groups in the variables “total time of moderate and vigorous PA” (p=.009 and p=0.0001, respectively) and “daily mean of moderate and vigorous physical activity” (p=0.005 and p=0.003), with patients performing worse than the control group (Table 2).

There was a statistically significant difference between groups in the “Metabolic Equivalent” (MET) variables, with p=0.04, “Total Steps” (p=0.04) and “total energy expenditure” (p=0.0001), with the lower values always occurring in the patients group (Table 2). In the studied sample, 8% of patients with SCA and 30% of controls complied with the current recommendations of PA, which is 60 minutes or more of moderate or vigorous PA (MVPA) per day [17].

There was no statistically significant difference in the average time of sedentary activity (neither in total or in the daily average), but these values were always lower in the control group.

The questionnaire (PAQ-C) was applied to all participants. The mean score obtained by the patients was 1.65 ± 0.4 and the mean score by the controls was 3.39 ± 0.38. 62% of the patients with SCA were categorised as very sedentary and the remaining 38% were sedentary (Table 3). Among the controls, 11% were classified as sedentary, 75% as moderately active, and 14% were active.

Table 3 presents the comparison of the PA level in the different activity categories evaluated by PAQ-C and shows that patients with SCA reported lower PA levels in all categories compared to healthy controls.

**Table 3:** Comparison of physical activity levels in different categories assessed by the PAQ-C between patients and controls.

| Variables | SCA children and adolescents (n=50) | Range | Controls (n=50) | Range | p |
| --- | --- | --- | --- | --- | --- |
| Spare-time activity | 0.75 ± 0.3 | 0 – 1.6 | 2.48 ± 0.61 | 1.4 – 3.7 | 0.0001 |
| Activity during PEC | 1.74 ± 0.6 | 1 – 3 | 3.36 ± 0.6 | 2 – 5 | 0.0001 |
| Break-time Activity | 1.6 ± 0.69 | 1 – 3 | 3.64 ± 0.59 | 2 – 5 | 0.0001 |
| Lunch-time Activity | 1.42 ± 0.53 | 1 – 3 | 3.32 ± 0.71 | 2 – 5 | 0.0001 |
| After school Activity | 1.8 ± 0.98 | 1 – 4 | 3.38 ± 0.56 | 2 – 4 | 0.0001 |
| Evening Activity | 1.7 ± 0.8 | 1 – 4 | 3.58 ± 0.7 | 2 – 5 | 0.0001 |
| Weekend Activity | 2.08 ± 0.75 | 1 – 3 | 3.63 ± 0.7 | 2 – 5 | 0.0001 |
| AF during the last 7days | 2.32 ± 0.81 | 1 – 4 | 3.72 ± 0.5 | 3 – 5 | 0.0001 |
| AF during each day last week | 1.62 ± 0.53 | 0.1 – 3 | 3.37 ± 0.58 | 2.5 – 4.3 | 0.0001 |
| Score total | 1.65 ± 0.4 | 0.8 – 2.3 | 3.39 ± 0.38 | 2.7 – 4.2 | 0.0001 |

PEC: Physical Education Classes; AF: Activity frequency; Data expressed as mean ± standard deviation and range (minimum and maximum). Independent t-test and Mann-Whitney U test were used to compare the two groups when the variables presented parametric and non-parametric distribution, respectively.

## Discussion

To the authors’ knowledge, this study presents the first results on sedentary time and the intensity level of PA of children and adolescents with SCA in Brazil. The results demonstrate increased sedentary behaviour and less intense PA levels in children and adolescents with SCA compared to healthy controls. These results are of particular importance when considering the beneficial effects on the oxidative stress damages, previously identified in a study which used an animal model [5]. The authors of that study proposed that an exercise program could be useful for controlling clinical complications due to SCA [5].

The effects of an exercise program applied to heterozygous carriers for HbS (sickle cell trait) were previously evaluated and a study reported beneficial results on endothelial function, including reduction of oxidative stress markers and antioxidant enhancement (increased activity and NO availability) [18]. There is evidence that sedentary behaviour is associated with adverse health effects in groups of individuals with various chronic diseases [19], but patients with SCA have never been evaluated until now.

Impairment of nutritional status and growth retardation in children and adolescents with SCA are associated with resting energy expenditure 10-20% higher than that observed in healthy individuals, which is at least partially due to the higher cardiac output, such as mechanisms of compensation for moderate or severe and chronic anemia [20]. The present study showed a statistically significant difference in the MET variable, which reinforces the findings of a previous study [21]. In order to maintain their total daily energy expenditure at the same level as healthy adolescents, patients with SCA reduce their energy expenditure in PA [22].

Previous studies have evaluated the activity of the autonomic nervous system in patients with SCA and have identified an imbalance caused by parasympathetic activity at rest [22] and deficiency of autonomic reactivity [23]. Furthermore, the degree of impairment is associated with clinical severity [24]. The present study did not evaluate the activity of the autonomic nervous system. However, in healthy individuals, the energy expended with PA is positively associated with the activity of the autonomic nervous system, especially the activity of the parasympathetic nervous system [25]. Regular PA increases the activity of the parasympathetic nervous system, which has a protective effect on the cardiovascular system [26], whilst sedentary behaviour leads to an imbalance in the autonomic nervous system activity, which may favour the development of cardiovascular diseases [27], a condition that is particularly detrimental to the patients with SCA.

The present study identified low PA levels and low energy expenditure in patients with SCA compared with healthy individuals, corroborating previous studies [19]. Various factors, such as muscular hypotrophy, pulmonary and cardiac complications, may explain these findings [5].

However, intense physical exercise induces metabolic and physiological changes that may be detrimental to individuals with SCA [5] and there is no consensus on the maximum intensity of safe exercise that these patients can tolerate. In addition, due to the limitations imposed by the disease and its frequent acute intercurrences, parents of children and adolescents with SCA may discourage them from engaging in physical activities [28], which may explain the low energy expenditure and physical activity in the sample studied.

A previous study identified positive effects in MVPA for patients with SCA [29] and considered this practice to be safe. Considering the findings of this study, future objectives are to identify which training modalities would be better tolerated and could provide the greatest health benefits to patients. Given the findings, it is suggested that the evaluation of PA should be part of the outpatient follow-up for patients with SCA, being an important tool to determine the severity of the disease and to suggest a possible strategy to prevent clinical complications.

We identified a limitation of the present study: the use of the accelerometer was voluntarily and in the absence of acute intercurrences. Thus, it is possible that patients with more severe forms of SCA have not been included. However, given the results obtained, it is assumed that the inclusion of patients with greater frequency or intensity of symptoms would result in less PA and a more sedentary lifestyle.

## Conclusion

Children and adolescents with SCA were assessed for PA, assessed subjectively by the PAC-C and objectively by the accelerometer, resulting in values lower than that of healthy children and adolescents.

The results indicate that programs with a focus on promoting optimal PA levels and on reducing sedentary behaviour in this population are necessary. These efforts must be intensified.

## Acknowledgments

The authors thank the Centre of Post-Graduation in Health Sciences of the Federal University of Sergipe for allowing the use of accelerometers, and to the assistance team of the Pediatric Hematology Unit of the University Hospital of the Federal University of Sergipe for making the patients’ clinical and laboratory data available. The authors are also grateful to the board of Pio X High School, to the parents of the students, for allowing the recruitment of healthy children and adolescents, and particularly to the patients with SCA who participated in the project and their parents.

